# Resistance is futile: A CRISPR homing gene drive targeting a haplolethal gene

**DOI:** 10.1101/651737

**Authors:** Jackson Champer, Emily Yang, Yoo Lim Lee, Jingxian Liu, Andrew G. Clark, Philipp W. Messer

## Abstract

Engineered gene drives are being explored as a potential strategy for the control of vector-borne diseases due to their ability to rapidly spread genetic modifications through a population. While an effective CRISPR homing gene drive for population suppression has recently been demonstrated in mosquitoes, formation of resistance alleles that prevent Cas9 cleavage remains the major obstacle for drive strategies aiming at population modification, rather than elimination. Here, we present a homing drive in *Drosophila melanogaster* that reduces resistance allele formation below detectable levels by targeting a haplolethal gene with two gRNAs while also providing a rescue allele. This is because any resistance alleles that form by end-joining repair will typically disrupt the haplolethal target gene, rendering the individuals carrying them nonviable. We demonstrate that our drive is highly efficient, with 91% of the progeny of drive heterozygotes inheriting the drive allele and with no resistance alleles observed in the remainder. In a large cage experiment, the drive allele successfully spread to all individuals. These results show that a haplolethal homing drive can be a highly effective tool for population modification.

## INTRODUCTION

The ability to modify the genomes of wild populations would provide a powerful new approach for reducing the transmission of vector-borne diseases such as malaria or dengue^1–4^. This could be achieved through engineered gene drives, which can spread rapidly through a population by biasing inheritance in their favor^1–7^ and thereby disseminate engineered payload alleles that manipulate an organism’s viability, fecundity, or ability to transmit disease.

Homing gene drives based on the CRISPR/Cas9 system have shown promise in several organisms including yeast^5–8^, flies^9–15^, mosquitoes^16–18^, and mice^19^. The mechanism of such homing drives involves RNA-guided Cas9 cleavage at a target site, allowing the drive allele to be copied to the wild-type chromosome in the germline through homology-directed repair. As a result, the drive allele will increase in frequency over time. However, Cas9-induced double-strand breaks can also be repaired through end-joining mechanisms, instead of homology-directed repair, which typically mutates the target sites. In most cases, this creates a resistance allele because the changed sequence at the target site will no longer be recognized by the guide RNA (gRNA) and thus be immune to further Cas9 cleavage. The formation of resistance alleles by this process has been observed to occur in the germline of drive-carrying organisms as well as in embryos due to maternally deposited Cas9^12^.

For drive strategies that aim to suppress a population, resistance alleles with a disrupted target site do not substantially impede the success of the drive. This is because suppression drives usually target an essential gene without providing rescue, thus allowing resistance alleles to contribute to the population suppression function of the drive, even if they are expected to somewhat retard the spread of the drive. By targeting a highly conserved site with an efficient Cas9 promoter that minimized resistance allele formation in both the germline and the embryo, a recent population suppression approach was indeed successful in eliminating small cage populations of *Anopheles gambiae*^20^. Nevertheless, it remains unclear how well such an approach would work in a larger population, where suppression drives face strong selection pressure for the evolution of resistance alleles that preserve target gene function. If even a small number of such alleles forms, this would be expected to quickly allow a population to rebound from the suppressive effects of a drive. Ecological factors may also prevent a population suppression drive from achieving complete success^21^. Thus, population modification strategies may still be desirable as alternatives to suppression strategies, particularly for applications such as reduction of malaria transmission for which suitable payload genes are already available^1^. However, the high rates at which resistance alleles typically form in population modification drives continues to pose a major hurdle for the successful spread of such drives.

One proposed strategy to overcome the problem of resistance in a modification drive is the haplolethal homing drive^22^. By targeting a haplolethal gene, any end-joining repair would be expected to produce lethal mutations, removing resistance alleles from the population as they form. However, the drive allele itself would contain a recoded portion of the haplolethal gene that cannot be cleaved by the drive and remains functional. This should eliminate individuals that inherit resistance alleles while allowing drive-carrying individuals to survive and reproduce normally. Here, we demonstrate that a haplolethal homing drive with two gRNAs can serve as an effective population modification system in *Drosophila melanogaster*. Our haplolethal drive construct had high rates of drive inheritance and no detectable formation of resistance alleles, which allowed it to successfully spread through a large cage population.

## METHODS

### Plasmid construction

The starting plasmids pCFD3^23^ (Addgene plasmid #49410), pCFD4^23^ (Addgene plasmid # 49411), and pCFD5^24^ (Addgene plasmid #73914) were kindly supplied by Simon Bullock. A starting plasmid similar to BHDcN1 was constructed in a previous study^13^ (see Supplemental Information). Restriction enzymes for plasmid digestion, Q5 Hot Start DNA Polymerase for PCR, and Assembly Master Mix for Gibson assembly were acquired from New England Biolabs. Oligonucleotides and gBlocks were obtained from Integrated DNA Technologies. JM109 competent cells and ZymoPure Midiprep kit from Zymo Research were used to transform and purify plasmids. A list of DNA fragments, plasmids, primers, and restriction enzymes used for cloning of each construct can be found in the Supplemental Information section.

### Generation of transgenic lines

Injections were conducted by Rainbow Transgenic Flies. The donor plasmid AHDr352v2 (744 ng/µL) was injected along with plasmid AHDrg2 (20 ng/µL), which provided additional gRNAs for transformation, and pBS-Hsp70-Cas9 (140 ng/µL, from Melissa Harrison & Kate O’Connor-Giles & Jill Wildonger, Addgene plasmid #45945) providing Cas9. A 10 mM Tris-HCl, 100 µM EDTA solution at pH 8.5 was used for the injection.

### Genotypes and phenotypes

Drive carriers are indicated by expression of dsRed drives by the 3×P3 promoter, which is highly visible in the eyes of *w*^*1118*^ flies. EGFP similarly marks flies carrying Cas9 driven by the *nanos* promoter^15^. For phenotyping, flies were anesthetized with CO_2_ and scored for red fluorescence in the eyes using the NIGHTSEA system (SFA-GR). Fly line homozygosity was assessed by fluorescence intensity and confirmed by sequencing.

**Fly rearing and phenotyping**

Flies were reared in Bloomington Standard medium and housed in an incubator at 25°C following a 14/10 hour day/night cycle. For the cage study, flies were housed in 30×30×30 cm (Bugdorm, BD43030D) enclosures. Initially, a fly line was generated that was homozygous for both the drive flies and the split-Cas9 allele. These, together with split-Cas9 homozygotes of the same age, were separately allowed to lay eggs in eight food bottles for a single day. Bottles were then placed in cages, and eleven days later, they were replaced in the cage with fresh food. Bottles were removed from the cages the following day, the flies were frozen for later phenotyping, and the egg-containing bottles returned to the cage. This 12-day cycle was repeated for each generation.

To minimize risk of accidental release, all live gene drive flies were quarantined at the Sarkaria Arthropod Research Laboratory at Cornell University under Arthropod Containment Level 2 protocols in accordance with USDA APHIS standards. In addition, the split-Cas9 drive system^15^ prevents drive conversion in wild-type flies, which lack the endonuclease. All safety standards were approved by the Cornell University Institutional Biosafety Committee.

### Genotyping

For genotyping, flies were frozen, and DNA was extracted by grinding flies in 30µL of 10 mM Tris-HCl pH 8, 1mM EDTA, 25 mM NaCl, and 200 µg/mL recombinant proteinase K (ThermoScientific), followed by incubation at 37°C for 30 minutes and then 95°C for 5 minutes. The DNA was used as a template for PCR to amplify the region of interest. After DNA fragments were isolated by gel electrophoresis, sequences were obtained by Sanger sequencing and analyzed with ApE software (http://biologylabs.utah.edu/jorgensen/wayned/ape).

## RESULTS

### Drive Dynamics

A CRISPR haplolethal homing gene drive contains all the elements of a standard homing drive, including a nuclease, gRNA, and an optional payload gene. It is located at the gRNA target site, and when the wild-type allele is cleaved in the germline, the break can be repaired by homology-directed repair, which usually results in successful copying (“homing”) of the drive (Figure 1). This converts the germline cell from a drive heterozygote into a homozygote. However, end-joining repair of the DNA cut can result in resistance allele formation, which can be of the “r1” type (preserving the function of any target gene) or the “r2” type (disrupting the target gene). The latter form is more common, since roughly two thirds of mutations will create frameshifts, and most of the remainder tend to still sufficiently change the amino acid sequence to disrupt protein function. In the progeny of females with at least one drive allele, additional resistance allele formation occurs in the offspring of females carrying the drive due to maternally deposited Cas9 and gRNAs. If cleavage occurs later in development of the embryo, mosaic resistance allele formation is possible, with different cells potentially containing a mixture of wild-type, r1, and r2 alleles^12,13^.

**FIGURE 1.**
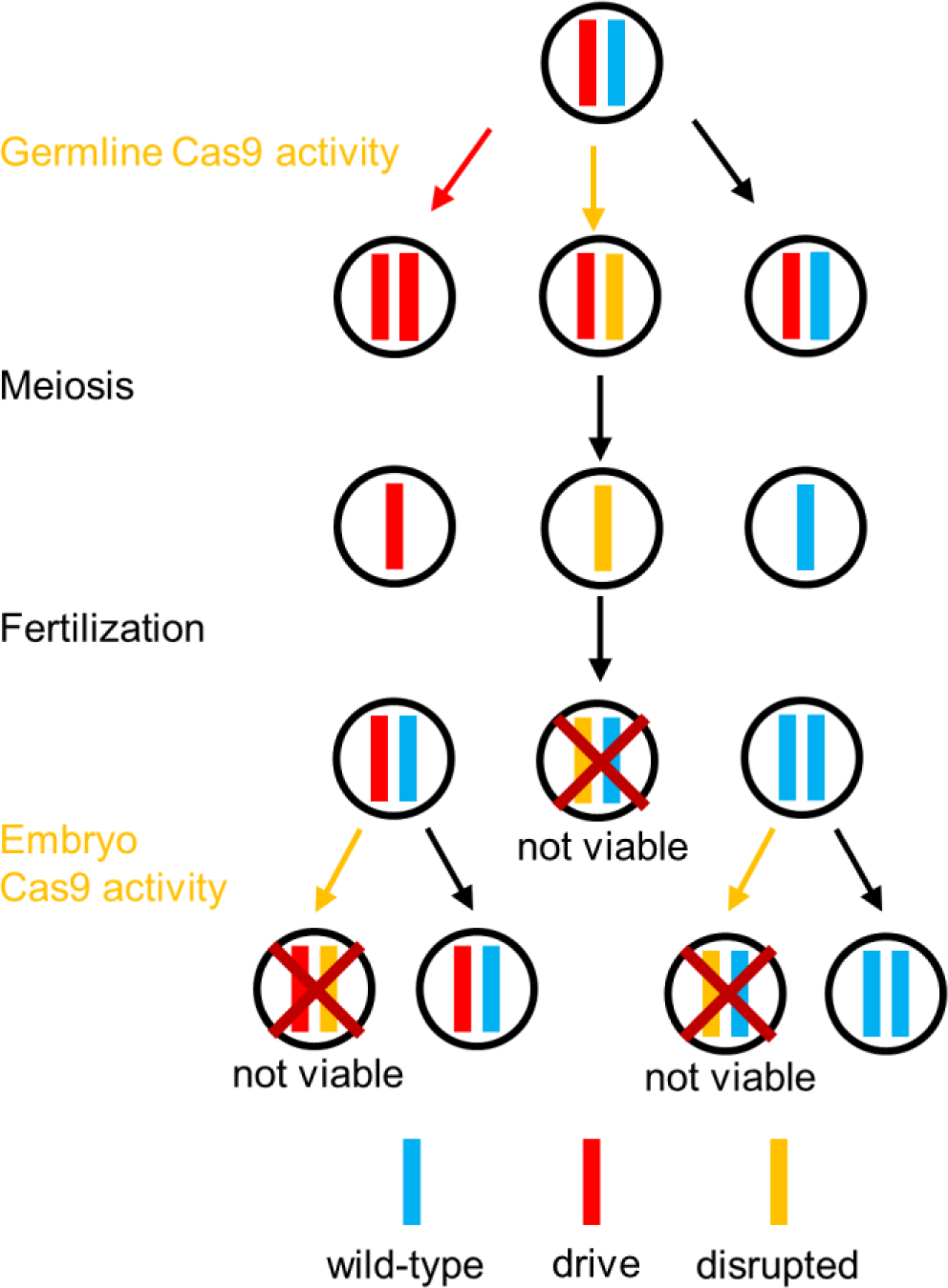
Haplolethal homing drive mechanism. Drive conversion occurs in the germline, though if Cas9 cleavage is repaired by end-joining, a resistance allele is formed. Additional resistance alleles can form due to maternally deposited Cas9 and gRNAs in the embryos from mothers that had at least one drive allele. All embryos that receive a resistance allele that disrupts the target gene will be nonviable.

In addition to containing the standard homing drive components, a haplolethal homing drive specifically targets a haplolethal gene, specifying a class of gene where two copies are needed for viability. The drive construct also carries a copy of the target gene to rescue gene function, which has been recoded with many synonymous sequence changes so that it is immune to Cas9 cleavage by the drive. However, if an r2 resistance allele is formed in either the embryo or in a germline precursor cell, the embryo will not survive, thereby eliminating such resistance alleles from the population. Embryos with high levels of mosaicism will likely also be nonviable, and lower levels may still impact fitness. Through this mechanism, the haplolethal homing drive largely negates the negative effects of forming resistance alleles in the germline, since these individuals will perish. However, resistance allele formation in the embryo will often result in drive-carrying individuals being rendered nonviable, so only a certain level can be tolerated before the drive would no longer be able to spread, with the specific critical thresholds being determined by the fitness costs of the drive.

### Drive construct design

Our drive construct (Figure 2) includes two gRNAs with targets 26 nucleotides apart in exon 4 (the second exon with a coding sequence) of the haplolethal gene *RpL35A*, a highly conserved protein component of the 60S ribosomal subunit^25^. Two tRNA sequences at the beginning and in between the gRNAs are endogenously spliced out from the pre-RNA^24^. The multiplexed gRNAs increase efficiency of the drive by providing two opportunities for cleavage, thus potentially increasing the rate of homology-directed repair and reducing the rate at which r1 alleles are formed^13^. The drive construct is flanked by two homology arms with wild-type sequences that abut the left and right gRNA target sites. Immediately adjacent to the left homology arm is a recoded version of the remainder of the coding sequence of *RpL35A*, followed by the wild-type 3’UTR. A dsRed sequence with the 3xP3 promoter produces red fluorescent protein expression in the eyes, allowing for easy identification of drive-carrying individuals. A split-drive element, containing Cas9 driven by the *nanos* promoter for expression in the germline and EGFP with the 3xP3 promoter, was previously constructed^15^ and is necessary for the drive to function, thus preventing it from spreading through wild-type individuals in the event of an accidental release^15^.

**FIGURE 2.**
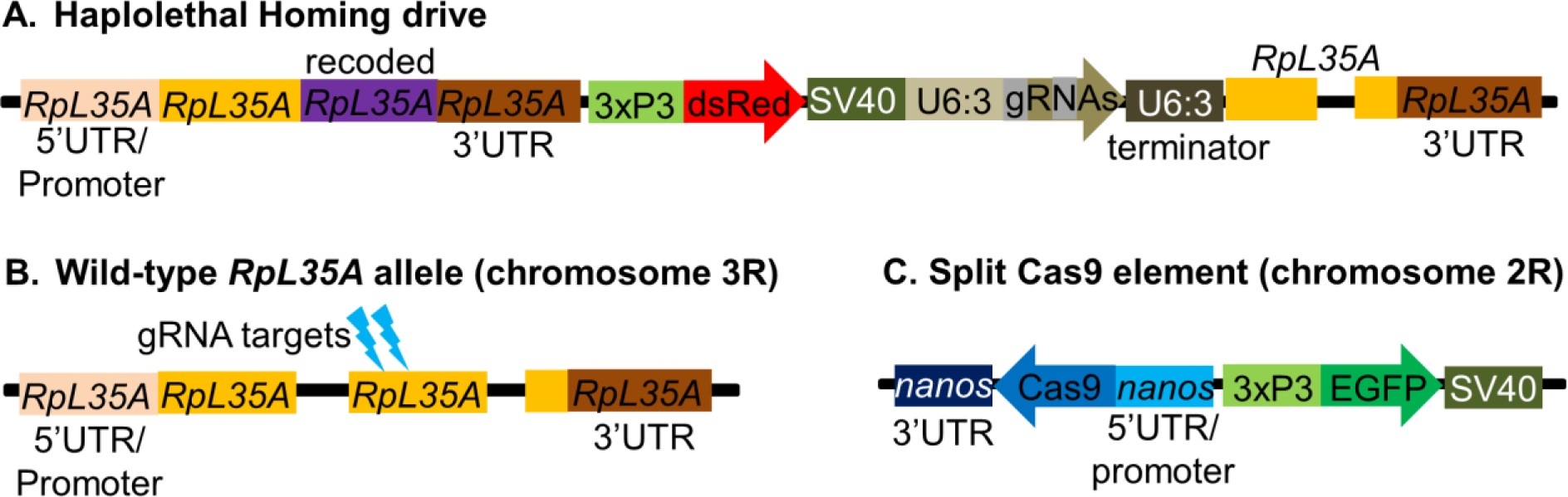
Drive Construct Schematic. (**A**) The drive construct contains a recoded version of *RpL35A*, dsRed, and gRNAs. (**B**) The wild-type allele of *RpL35A* is cleaved in the second coding exon by the drives two gRNAs. (**C**) The supporting element of the drive contains Cas9 with the *nanos* promoter and 3’UTR with an EGFP marker.

### Drive inheritance

Flies homozygous for the drive were crossed with a laboratory line homozygous for the *nanos*-Cas9 allele to produce offspring that were heterozygous for the drive and split-Cas9 element. These individuals were then crossed to *w*^*1118*^ flies to assess drive conversion efficiency. Observation of the dsRed phenotype in the eyes of the offspring from this cross was used to identify drive carriers. The progeny of both heterozygous females and of heterozygous males showed 91% drive inheritance (Figure 3, Data Set S1-S2), which were both significantly different from the 50% expected under Mendelian inheritance (*p*<0.0001, Fisher’s exact test). To assess formation of resistance alleles, seventeen offspring from the cross between drive females and *w*^*1118*^ males were sequenced at the target locus. Sixteen (eleven of which also had a drive allele) were found to be completely wild type, and one (that also had a drive allele) was found to be mosaic, indicating that few or no flies possessed viable resistance alleles. This is in stark contrast to previous homing drives in *D. melanogaster*, which all had high resistance rates, thus implying that resistance alleles for the haplolethal homing drive were all of the r2 type, and flies inheriting them were all nonviable. The mosaicism seen in one individual was likely produced by embryo Cas9 activity later during development, creating a resistance allele without enough penetrance to be lethal. Such a resistance allele would presumably still become lethal if it were transmitted to progeny in a subsequent generation.

**FIGURE 3.**
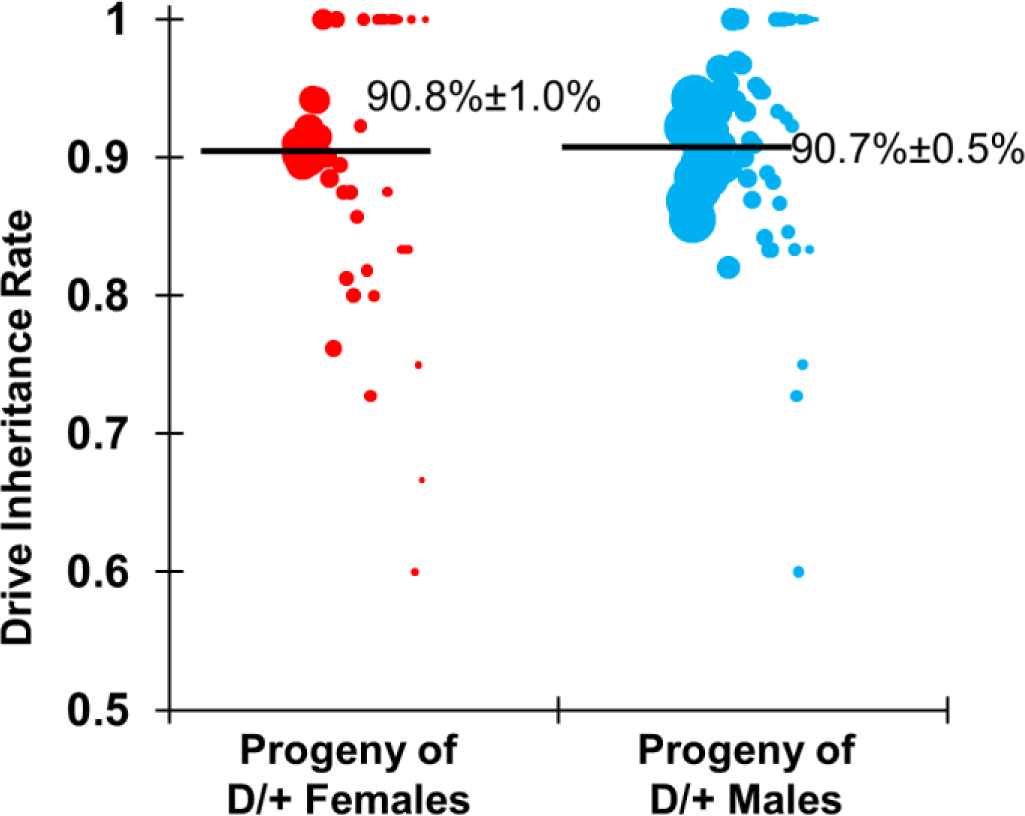
Drive inheritance rate. Flies heterozygous for the drive and Cas9 alleles were crossed with *w*^*1118*^ flies. The progeny of these were evaluated for drive inheritance as a percentage of flies with dsRed phenotype. The size of each dot represents the clutch size of one individual. The mean and standard deviation are indicated.

### Drive performance analysis

To gain a better understanding of the performance of the drive, individual females or males with one copy of the drive and one copy of the Cas9 allele were crossed with single *w*^*1118*^ flies, and *w*^*1118*^ flies were separately crossed together. Flies were allowed to lay eggs for a day, which were counted, and then pupae were counted seven days later to assess egg viability. The progeny of drive heterozygous males showed a pupae-to-egg survival rate of 80% (Figure 4, Data Set S2), which was similar to the 83% survival of eggs from *w*^*1118*^ flies (Data Set S3). The progeny of females had a pupae-to-egg survival of 55% (Figure 4, Data Set S1), a significant deviation from the viability of eggs from *w*^*1118*^ flies (*p*<0.0001, Fisher’s exact test). This is consistent with expectations that progeny carrying a resistance allele are nonviable and that resistance alleles form in the offspring of drive-carrying females due cleavage activity in the embryo from maternally deposited Cas9.

**FIGURE 4.**
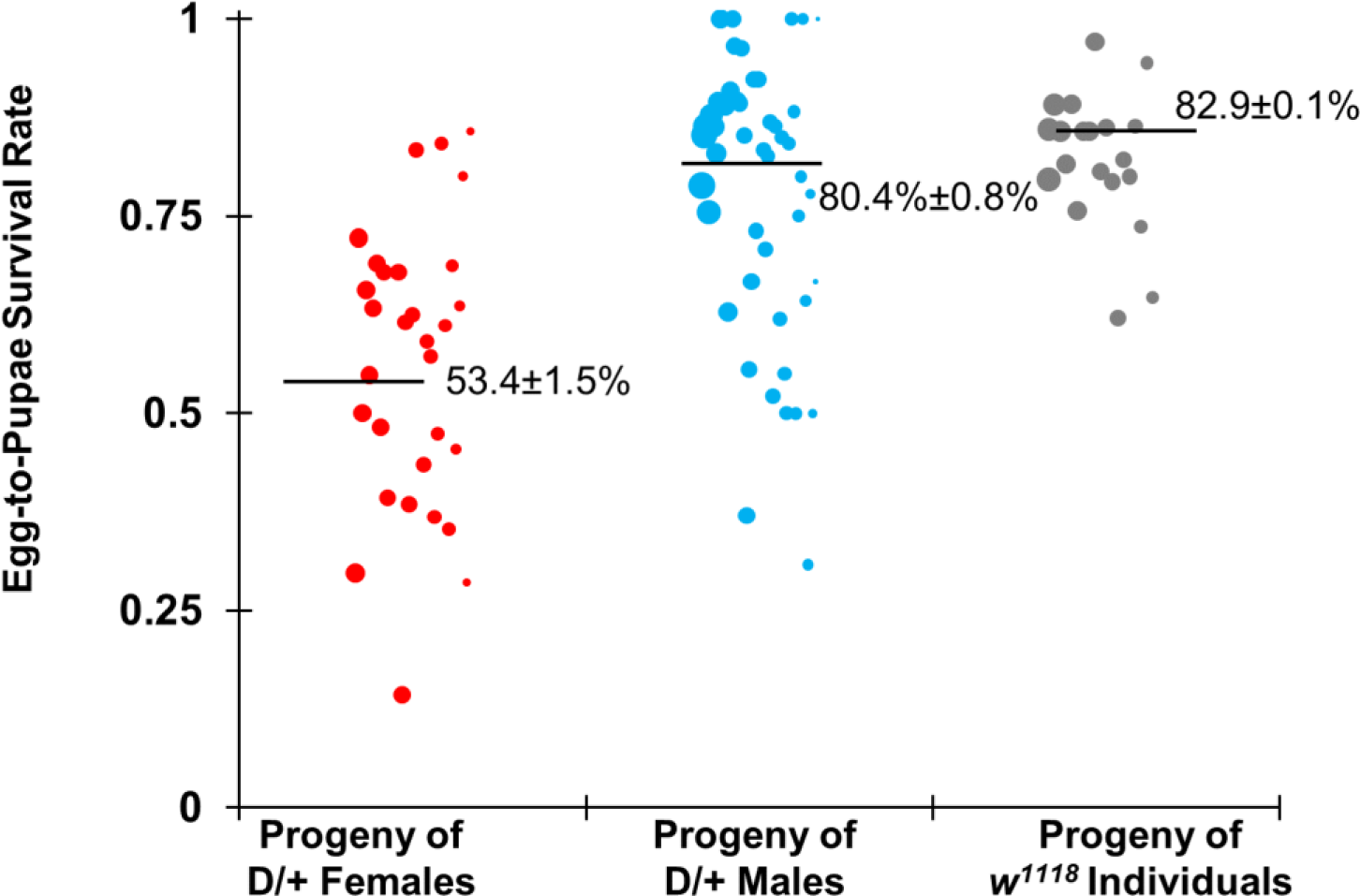
Pupae-to-egg survival rate. Females and males heterozygous for the drive and Cas9 alleles were crossed with *w*^*1118*^ flies, and *w*^*1118*^ individuals were crossed together. Females were allowed to lay eggs for one day, which were counted and then assessed for survival to the pupal stage. The progeny of heterozygous females had a significantly lower survival rate than wild-type flies, but the progeny of heterozygous males did not, with the difference presumably caused by reduced viability from resistance alleles that formed in the embryo due to maternally deposited Cas9 and gRNA. The size of each dot represents the clutch size of one individual. The means and standard deviations are indicated.

From viability data based on male crosses, germline resistance allele formation was likely low, at approximately 6% (Data S2). Based on the phenotypes of surviving progeny, this yields a drive conversion rate of 76%, with 18% of wild-type alleles remaining unconverted to drive or resistance alleles. The number of remaining wild-type individuals is substantially higher than in previous studies^12,13,15^, including one where we utilized the same split-Cas9 element^15^. This implies that either gRNA expression in the haplolethal homing drive is low or that the gRNAs have lower activity. Nevertheless, the drive conversion rate in males was higher than observed for previous drives in *D. melanogaster*, likely due to the multiplexed gRNAs. Because of this, a germline resistance allele formation rate of 6% may be an underestimate, since the rate of drive conversion appears to be similar to expected values based on the rates from previous studies^12,13,15^, but the resistance formation rate is lower. The relatively high error (4%) in the estimate of this resistance rate would be consistent with this notion.

In previous *D. melanogaster* drives, females had somewhat higher drive conversion rates than males, but also, with the exception of a drive targeting *yellow*, very high embryo resistance rates. The embryo resistance rates seen in our haplolethal homing drive were substantially lower, with two thirds of offspring surviving relative to wild-type crosses. This further supports the idea that the gRNA expression or activity in this particular drive is lower than that in previous drives. Additionally, nonviable eggs from drive females could be the result of both germline and embryo activity, which causes difficulties in disentangling the individual rates. One possible set of parameters that is consistent with the egg viability rate is a drive conversion rate of 76%, a resistance allele formation rate of 9%, and an embryo resistance rate of 28% (Data Set S1).

### Cage study

In a standard haplolethal homing drive with integrated Cas9, the Cas9 gene would be copied in most gametocytes, bringing the embryo Cas9 cleavage rates closer to the rate found in the embryos of individuals that are homozygous for the drive. To measure the effects of increased Cas9 expression, homozygotes for both the drive and split-Cas9 allele were generated. These were then crossed to Cas9 homozygotes with no drive, generating individuals that were drive/wild-type heterozygotes at the drive locus with two copies of Cas9. When such flies were crossed to *w*^*1118*^ individuals, both female and male drive heterozygotes showed 93% drive inheritance, which was slightly higher than for individuals with one copy of Cas9 (though this did not reach statistical significance). Since the additional Cas9 allele is expected to result in slightly higher germline cut rates due to more Cas9 expression, we approximated drive performance for the cage study in both males and females to be 78% drive conversion rate, 10% germline resistance allele formation rate, and 28% embryo resistance allele formation rate in the progeny of females.

To assess the performance of the haplolethal homing drive in large cage populations, flies homozygous for both the drive and Cas9 were allowed to lay eggs in bottles for one day, and flies homozygous for the Cas9 allele were similarly allowed to lay eggs in separate bottles. Flies were then removed, and the bottles were placed together in a population cage. The cage was followed for several generations, and each generation was phenotyped for dsRed to measure the frequency of drive carriers, which includes both homozygotes and heterozygotes (Figure 5, Data Set S4). The drive carrier frequency rose from 32% in generation zero (all of which were drive homozygotes) to 100% in generation six. In the final generation, approximately 1% of flies had a fainter dsRed phenotype, indicating that they potentially had only one drive allele. Fifteen of these individuals were genotyped, and it was found that ten were actually drive homozygotes while five had a remaining wild-type allele. This small number of wild-type alleles would likely have been eliminated or converted to drive alleles within one or two additional generations.

**FIGURE 5.**
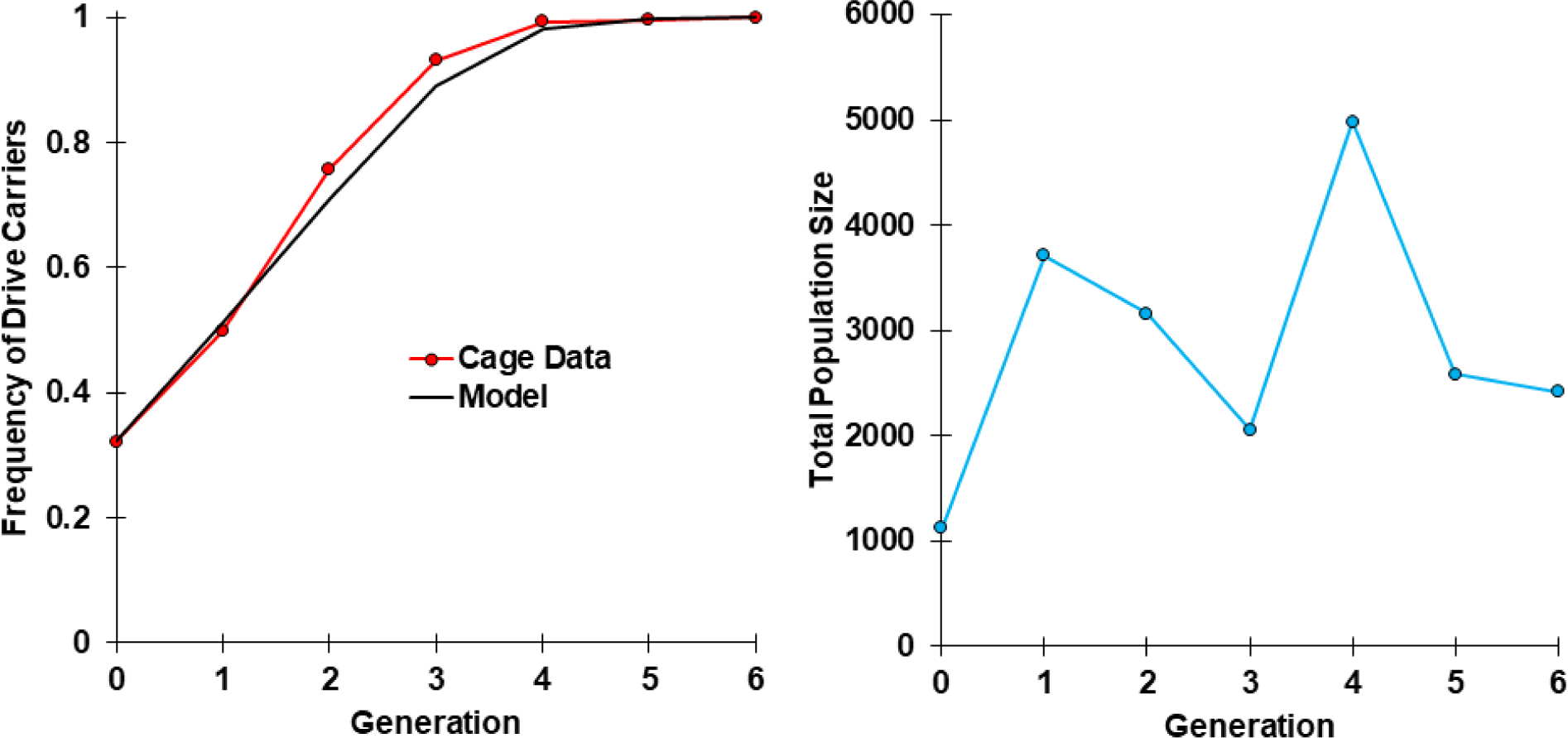
Cage study. Homozygotes for the drive and Cas9 alleles were introduced into a cage containing homozygotes for the Cas9 allele at an initial frequency of 32%, and the cage was followed for several discrete generations. All individuals from each generation were phenotyped for dsRed, indicating the presence of one or two drive alleles. The model represents a population with no fitness costs, a homing rate of 78%, a germline resistance allele formation rate of 10%, and an embryo resistance allele formation rate of 28%.

To assess the fitness of the drive allele, we adapted a maximum likelihood approach we developed previously^26^ for gene drive inheritance and a haplolethal target. Drive performance for both males and females with two copies of Cas9 was estimated as above. Separate cleavage at each gRNA was not modeled, but to take into account the low rate of r1 formation, all embryos with resistance alleles were assumed to be nonviable. A single fitness parameter was inferred for drive homozygotes that represented the relative fecundity for females and relative mating success for males compared to wild-type individuals. Drive heterozygotes were assigned fitness values equal to the square root of the value for drive homozygotes, equivalent to a co-dominant model. Because drive homozygotes and heterozygotes could not be accurately distinguished, they were assumed to be in relative proportions predicted by the model. With these parameters, the effective population size (set to be the average of the census size of the two generations in each generational transition) of the cage was inferred to be 5.6% of the census population size, with a 95% confidence interval of 1.6% to 13.8%. This estimate was similar to previous cages^26^, but the wider confidence interval was likely due to our inability to distinguish between drive homozygotes and heterozygotes. Our estimate of the fitness parameter was 1.1, indicating a higher fitness than wild-type individuals. However, the 95% confidence interval extended from 0.90 to 1.34. Thus, it is likely that drive fitness is similar to wild-type alleles. This implies that if rare r1 resistance alleles form as the drive sweeps through a population, it would likely be an extended period of time before such r1 alleles reach an appreciable number of individuals.

## DISCUSSION

We have demonstrated that a homing gene drive targeting a haplolethal gene can successfully eliminate resistance alleles while preserving drive carriers by including a rescue allele. This mechanism enabled our drive to rapidly spread to all individuals in a population cage, thus overcoming the key obstacle that has prevented success in previous homing drives designed for population modification.

Drives based on this mechanism could be used for rapid modification of entire populations with lower risk of disrupting the surrounding ecosystems than suppression drives. Population modification drives are also expected to be less vulnerable to complex ecological factors that may prevent the complete success of a suppression drive^21^, suggesting their potential use in conjunction with such drives. Many proposed applications for modification drives, such as the spreading of a disease-refractory payload allele through an insect vector population, should also still be able to function successfully in the face of a small level of resistance allele formation, which is unlikely to be the case for population suppression approaches. Furthermore, a variety of possible payload genes have already been demonstrated, including for the reduction of dengue and malaria transmission in *Aedes* and *Anopheles*, respectively^1^.

While germline resistance alleles are easily eliminated by a haplolethal homing drive without apparent negative effects on its performance, embryo resistance remains a major obstacle. Indeed, the high embryo resistance rates seen in previous *D. melanogaster* constructs^12,13,15^ would have prevented the successful spread of the drive. This is because in the haplolethal homing system, Cas9 activity in the embryo renders such embryos nonviable, most of which would possess a drive allele. Reducing embryo resistance, for example through the use of different promoters for Cas9, would be expected to greatly increase the overall efficiency of the drive. A similar consideration is somatic expression, where resistance alleles could impose a substantial fitness cost on drive/wild-type heterozygotes by disrupting the wild-type alleles in many cells. Although the *nanos* promoter utilized in this study has minimal leaky somatic expression, somatic expression may be an important consideration for other promoters. This potentially includes the *zpg* promoter used successfully in an *Anopheles gambiae* population suppression drive, which appears to have a fecundity phenotype consistent with a small amount of somatic expression^20^.

Incomplete homology-directed repair could also pose a problem for a haplolethal homing drive if the recoded rescue allele is copied but other elements (particularly the payload) are not. Such partial homology-directed repair has been observed before in homing drives^10,12,13^, but it is unclear how often a large enough portion to provide rescue would be copied, while not copying the payload. One potential solution to this issue would be to use a “distant site” haplolethal homing drive. In this method, a full recoded rescue allele of the haplolethal gene with promoter would be included within a gene drive, which would be placed in a different location from the haplolethal gene. The drive would possess gRNAs for cleavage both at its own site for homology-directed repair and for disruption of the haplolethal target gene. Disrupted haplolethal gene alleles would therefore be paired with germline resistance alleles and drive alleles in the next generation, resulting in the former of these being nonviable. The drive alleles, due to their rescue element, would remain viable. This would largely avoid partial homology-directed repair of the rescue element if the payload and rescue alleles were both near the middle of the drive, yet drive performance may still suffer due to the presence of gRNAs not contributing to the copying of the drive and because of the increased size of the drive to accommodate a full rescue allele. Drives targeting haplolethal genes could also be redesigned to operate without homology-directed repair for population modification or suppression, though such drives would likely have substantially different dynamics^27^. Similar drives targeting recessive lethal genes have already been constructed^28,29^.

Although the fitness of our drive was indistinguishable from that of the wild-type allele in our model, it is possible that the model itself could be improved. For example, it is unclear if resistance alleles formed in the germline of males and possibly females are eliminated by the gamete stage or only later in the embryo, as we modeled it. If eliminated earlier, overall drive efficiency would be improved, which possibly could explain the slightly better-than-expected performance we saw in the cage experiment.

Our study demonstrates the efficiency of a multiple gRNA homing drive targeting a haplolethal gene with rescue. Since haplolethal genes are fairly widespread, the selection of targets would presumably be straightforward in many potential target species. While the need for a germline-specific promoter with minimal maternal carryover remains a prerequisite for the development of such as drive, we have shown that a less effective promoter can potentially still be viable if gRNA expression or activity is low. Future studies should test the implementation of our drive system into other organisms, including mosquitos. Large cage studies should further assess the r1 formation rate from partial homology-directed repair and the fitness cost of the drive, which will be critical for accurate modeling of the effectiveness of population modification strategies employing such drives.

## ACKNOWLEDGEMENTS

Thanks to Yassi Hafezi for helpful advice on the experimental work. This study was supported by the National Institutes of Health award R21AI130635 to J.C., A.G.C., and P.W.M, and the National Institutes of Health award F32AI138476 to J.C.

## SUPPLEMENTAL INFORMATION

|  |  |  |  |
| --- | --- | --- | --- |
| **AHDr352v2i1** | *Template* | *Oligo/Enzyme 1* | *Oligo/Enzyme 2* |
| *PCR Product* | gRpL35AG2 | RpL35A2g_F | RpL35A2g_R |
| *Plasmid Digest* | pCFD4 | BbsI | none |
| **AHDr35Ni1** | *Template* | *Oligo/Enzyme 1* | *Oligo/Enzyme 2* |
| *PCR Product* | Genomic DNA | R35Left_F | R35Left_R |
| *PCR Product* | gR35Left | gR35Left_F | gBlockLeft_R |
| *Plasmid Digest* | BHDrN1* | KpnI | XbaI |

\*Similar to previously constructed plasmid BHDrN1<sup>13</sup> for purposes of cloning in this study, but with the following sequence immediately 5' of the ApaI site sequence:

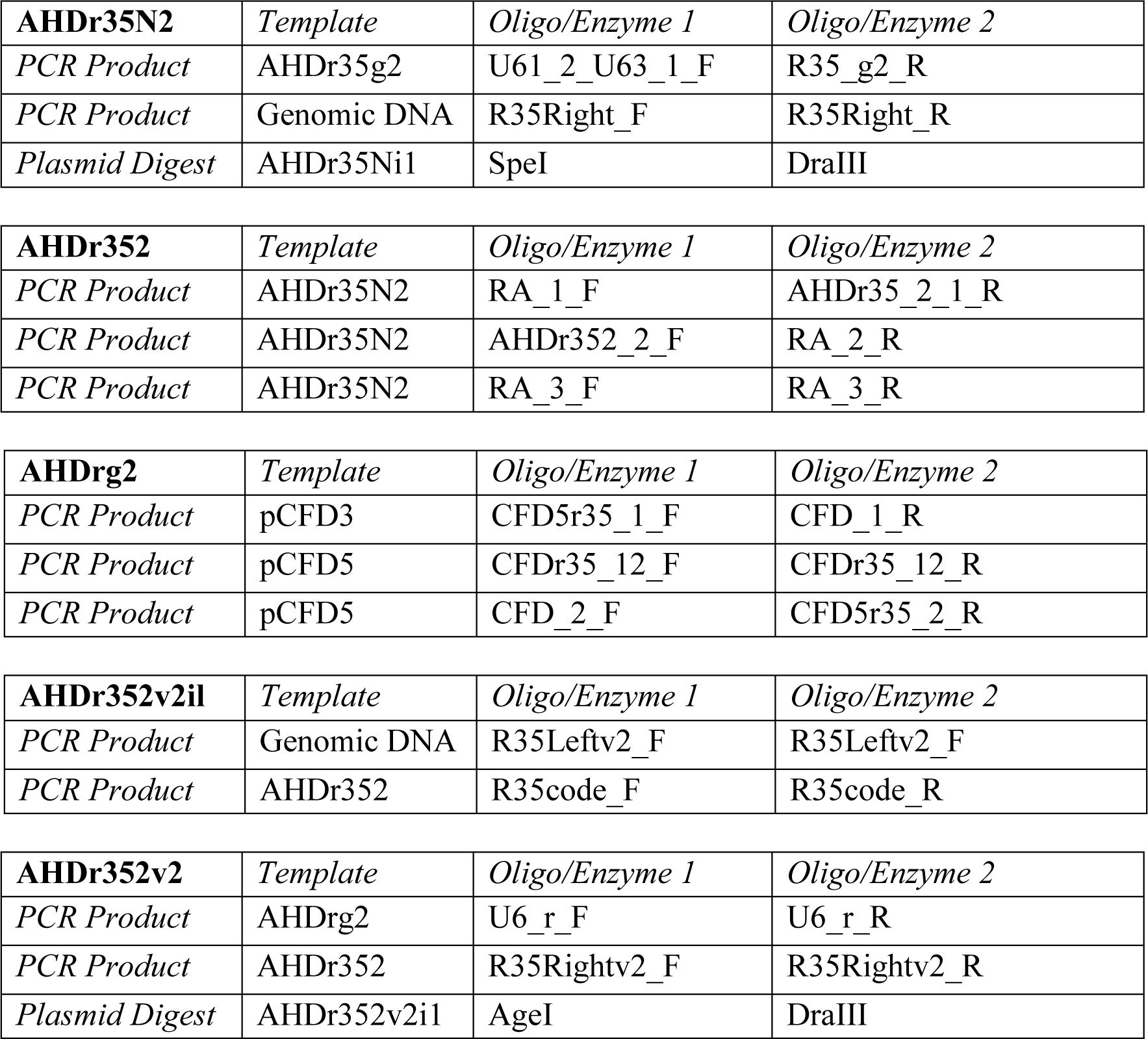

### Construction primers

~~~
AHDr35_2_1_R: TGAATTAGATCCCGGGAGTAGGGAAAGTCAAACCGAA
AHDr352_2_F: TGACTTTCCCTACTCCCGGGATCTAATTCAATTAGAGACTAATTCAA
CFD_1_R: GGCTATGCGTTGTTTGTTCTGC
CFD_2_F: AACAGTAGGCAGAACAAACAACGC
CFD5r35_1_F: GTGCAGCGTTGCGTCTATGTCTACAGTTTTAGAGCTAGAAATAGCAAGTTAAA
CFD5r35_2_R: AAAACACGTAGAAGGATCCGTGCTCTGCATCGGCCGGGAATCGAAC
CFDr35_12_F: ATGCAGAGCACGGATCCTTCTACGTGTTTTAGAGCTAGAAATAGCAAGTTAAA
CFDr35_12_R: AAAACTGTAGACATAGACGCAACGCTGCACCAGCCGGGAATCGAAC
gBlockLeft_R: CGAAAAGGGCCAGGAAGGAGCA
gR35Left_F: AGGCGCCTAAGGCCGAGAAGC
R35_g2_R: TGGTCTCGGCCTTGTATGCATACGCATTAAGCGAACA
R35code_F: GCACGGATCCTTCTATGTGGGCAAGCGCTGCGT
R35code_R: AATTGAATTAGTCTCTAATTGAATTAGATCCCGGGAGTAGGGAAAGTCAAACCGAACAGC
R35Left_F: ATTAACCAATTCTGAACATTATCGCCTAGGGTACCAACAGCACACTTTCGAGCAACGGCG
R35Left_R: AGGCGGCGGGCTTCTCGGCCTTAG
R35Leftv2_F: ACCAATTAACCAATTCTGAACATTATCGCCTAGGGTACCAACAGCACACTTTCGAGCAAC
R35Leftv2_R: CAGCGCTTGCCCACATAGAAGGATCCGTGCTCCTTG
R35Right_F: TTAATGCGTATGCATACAAGGCCGAGACCAAGAAGTGC
R35Right_R: GACGGAAGAGCCTCGAGCTGCACACACAGTGTCGGCTATAATTCTGACACATACCAAATG
R35Rightv2_F: TTAATGCGTATGCATACAAGGCCGAGACCAAGAAG
R35Rightv2_R: GATTGACGGAAGAGCCTCGAGCTGCACACACAGTGTCGGCTATAATTCTGACACATACCA
RA_1_F: ATTTATCAGCAATAAACCAGCCA
RA_2_R: GAACAACTCTCAGGCTCCAG
RA_3_F: TTTTGCCTACCTGGAGCCTGA
RA_3_R: TTCCGGCTGGCTGGTTTATTG
RpL35A2g_F: GGAAAGATATCCGGGTGAACTTCGCGTTGC
RpL35A2g_R: CTTGCTATTTCTAGCTCTAAAACCCTCAGAGGCTAA
U6_r_F: TTGTCCAAACTCATCAATGTATCTTAACCGGTGAGCTCTTTTTTGCTCACCTGTGATTGC
U6_r_R: TGGTCTCGGCCTTGTATGCATACGCATTAAGCGAACA
U61_2_U63_1_F: GTATGCTATACGAAGTTATAGAAGAGCACTAGTATTTTCAACGTCCTCGATAGTATAGT
~~~

### Sequencing Primers

~~~
pCFD5_S_R: ACGTCAACGGAAAACCATTGTCTA
RpL35ALeft_S_F: GCATGCAAATGATCGAAACCCT
RpL35ALeft_S_R: AGTGGACTTGGCTTGTGTGTC
RpL35ARight_S_F: CGCATCCGCATCGTTAGTTCA
RpL35ARight_S_R: TGCAGGTCAGTAATTCAAGTCGG
RpL35ARight_S2_R: CGTTTCCATCGTCTTCATCTGC
~~~

### gBlocks

~~~
gR35Left:
AGGCGCCTAAGGCCGAGAAGCCCGCCGCCTCCGAGGCCAAGGTGAGCGCCAAGAAGTATAAGCGCCATGGCCGCCTG TTTGCCAAGGCCGTGTTTACGGGATATAAGCGCGGCCTGCGCAATCAGCATGAGAATCAGGCCATTCTGAAGATCGA GGGAGCCCGCCGCAAGGAGCATGGCAGCTTTTATGTGGGCAAGCGCTGCGTGTACGTGTATAAGGCCGAGACGAAGA AGTGCGTCCCCCAGCACCCGGAGCGCAAGACGCGCGTGCGCGCCGTGTGGGGAAAGGTGACGCGCATTCATGGAAAT ACGGGAGCCGTCCGCGCCCGCTTTAATCGCAATCTGCCGGGCCACGCCATGGGACATCGCATTCGCATTATGCTGTA TCCCAGCCGCATCTAAGTTAATATCCGACTTGAATTACTGACCTGCAGGAGTAAAAAATCCGTTTTACATTAAATGA AACACTTTAAATTTAATTAAAACGCAACTTGGCTTTTTTATTAAGGCGAGATACCGATTGAAAGTTGACGGTAATCT GTATATCGATTGATGGCTGTTCGGTTTGACTTTCCCTACTTCTAGACATGCTCCTTCCTGGCCCTTTTCGA
gRpL35AG2:
GGAAAGATATCCGGGTGAACTTCGCGTTGCGTCTATGTCTACAGTTTTAGAGCTAGAAATAGCAAGTTAAAATAAGG CTAGTCCGTTATCAACTTGAAAAAGTGGCACCGAGTCGGTGCTTTTTTGCTCACCTGTGATTGCTCCTACTCAAATA CAAAAACATCAAATTTTCTGTCAATAAAGCATATTTATTTATATTTATTTTACAGGAAAGAATTCCTTTTAAAGTGT ATTTTAACCTATAATGAAAAACGATTAAAAAAAATACATAAAATAATTCGAAAATTTTTGAATAGCCCAGGTTGATA AAAATTCATTTCATACGTTTTATAACTTATGCCCCTAAGTATTTTTTGACCATAGTGTTTCAATTCTACATTAATTT TACAGAGTAGAATGAAACGCCACCTACTCAGCCAAGAGGCGAAAAGGTTAGCTCGCCAAGCAGAGAGGGCGCCAGTG CTCACTACTTTTTATAATTCTCAACTTCTTTTTCCAGACTCAGTTCGTATATATAGACCTATTTTCAATTTAACGTC GGAAACTTTAGCCTCTGAGGGTTTTAGAGCTAGAAATAGCAAG
~~~

## REFERENCES

1. Champer J, Buchman A, Akbari OS. Cheating evolution: engineering gene drives to manipulate the fate of wild populations. Nat Rev Genet, 17, 146–159, 2016.

2. Burt A. Heritable strategies for controlling insect vectors of disease. Philos Trans R Soc L B Biol Sci, 369, 20130432, 2014.

3. Alphey L. Genetic control of mosquitoes. Annu Rev Entomol, 59, 205–224, 2014.

4. Esvelt KM, Smidler AL, Catteruccia F, Church GM. Concerning RNA-guided gene drives for the alteration of wild populations. Elife, e03401, 2014.

5. DiCarlo JE, Chavez A, Dietz SL, Esvelt KM, Church GM. Safeguarding CRISPR-Cas9 gene drives in yeast. Nat Biotechnol, 33, 1250–1255, 2015.

6. Roggenkamp E, Giersch RM, Schrock MN, Turnquist E, Halloran M, Finnigan GC. Tuning CRISPR-Cas9 gene drives in Saccharomyces cerevisiae. G3, 8, g3.300557.2017, 2018.

7. Basgall EM, Goetting SC, Goeckel ME, Giersch RM, Roggenkamp E, Schrock MN, Halloran M, Finnigan GC. Gene drive inhibition by the anti-CRISPR proteins AcrIIA2 and AcrIIA4 in Saccharomyces cerevisiae. Microbiology, 2018.

8. Shapiro RS, Chavez A, Porter CBM, Hamblin M, Kaas CS, DiCarlo JE, Zeng G, Xu X, Revtovich A V., Kirienko N V., Wang Y, Church GM, Collins JJ. A CRISPR–Cas9-based gene drive platform for genetic interaction analysis in Candida albicans. Nat Microbiol, 3, 73–82, 2018.

9. Oberhofer G, Ivy T, Hay BA. Behavior of homing endonuclease gene drives targeting genes required for viability or female fertility with multiplexed guide RNAs. Proc Natl Acad Sci, 115, E9343–E9352, 2018.

10. KaramiNejadRanjbar M, Eckermann KN, Ahmed HMM, Sánchez C. HM, Dippel S, Marshall JM, Wimmer EA. Consequences of resistance evolution in a Cas9-based sex-conversion suppression gene drive for insect pest management. Proc Natl Acad Sci, 201713825, 2018.

11. Gantz VM, Bier E. Genome editing. The mutagenic chain reaction: a method for converting heterozygous to homozygous mutations. Science (80-), 348, 442–444, 2015.

12. Champer J, Reeves R, Oh SY, Liu C, Liu J, Clark AG, Messer PW. Novel CRISPR/Cas9 gene drive constructs reveal insights into mechanisms of resistance allele formation and drive efficiency in genetically diverse populations. PLoS Genet, 13, e1006796, 2017.

13. Champer J, Liu J, Oh SY, Reeves R, Luthra A, Oakes N, Clark AG, Messer PW. Reducing resistance allele formation in CRISPR gene drive. Proc Natl Acad Sci, 115, 5522–5527, 2018.

14. Champer J, Wen Z, Luthra A, Reeves R, Chung J, Liu C, Lee YL, Liu J, Yang E, Messer PW, Clark AG. CRISPR gene drive efficiency and resistance rate is highly heritable with no common genetic loci of large effect. Genetics, 2019.

15. Champer J, Chung J, Lee YL, Liu C, Yang E, Wen Z, Clark AG, Messer PW. Molecular safeguarding of CRISPR gene drive experiments. Elife, 8, 2019.

16. Hammond AM, Kyrou K, Bruttini M, North A, Galizi R, Karlsson X, Kranjc N, Carpi FM, D’Aurizio R, Crisanti A, Nolan T. The creation and selection of mutations resistant to a gene drive over multiple generations in the malaria mosquito. PLOS Genet, 13, e1007039, 2017.

17. Hammond A, Galizi R, Kyrou K, Simoni A, Siniscalchi C, Katsanos D, Gribble M, Baker D, Marois E, Russell S, Burt A, Windbichler N, Crisanti A, Nolan T. A CRISPR-Cas9 gene drive system targeting female reproduction in the malaria mosquito vector Anopheles gambiae. Nat Biotechnol, 34, 78–83, 2015.

18. Gantz VM, Jasinskiene N, Tatarenkova O, Fazekas A, Macias VM, Bier E, James AA. Highly efficient Cas9-mediated gene drive for population modification of the malaria vector mosquito Anopheles stephensi. Proc Natl Acad Sci U S A, 112, E6736–E6743, 2015.

19. Grunwald HA, Gantz VM, Poplawski G, Xu X-RS, Bier E, Cooper KL. Super-Mendelian inheritance mediated by CRISPR–Cas9 in the female mouse germline. Nature, 566, 105–109, 2019.

20. Kyrou K, Hammond AM, Galizi R, Kranjc N, Burt A, Beaghton AK, Nolan T, Crisanti A. A CRISPR-Cas9 gene drive targeting doublesex causes complete population suppression in caged Anopheles gambiae mosquitoes. Nat Biotechnol, 2018.

21. North AR, Burt A, Godfray HCJ. Modelling the potential of genetic control of malaria mosquitoes at national scale. BMC Biol, 17, 26, 2019.

22. Noble C, Olejarz J, Esvelt K, Church G, Nowak M. Evolutionary dynamics of CRISPR gene drives. Sci Adv, 3, e1601964, 2017.

23. Port F, Chen HM, Lee T, Bullock SL. Optimized CRISPR/Cas tools for efficient germline and somatic genome engineering in Drosophila. Proc Natl Acad Sci U S A, 111, E2967–76, 2014.

24. Port F, Bullock SL. Augmenting CRISPR applications in Drosophila with tRNA-flanked sgRNAs. Nat Methods, 13, 852–854, 2016.

25. Marygold SJ, Roote J, Reuter G, Lambertsson A, Ashburner M, Millburn GH, Harrison PM, Yu Z, Kenmochi N, Kaufman TC, Leevers SJ, Cook KR. The ribosomal protein genes and Minute loci of Drosophila melanogaster. Genome Biol, 8, R216, 2007.

26. Liu J, Champer J, Langmüller AM, Liu C, Chung J, Reeves R, Luthra A, Lee YL, Vaughn AH, Clark AG, Messer PW. Maximum Likelihood Estimation of Fitness Components in Experimental Evolution. Genetics, 211, 1005–1017, 2019.

27. Champer J, Kim I, Champer SE, Clark AG, Messer PW. Performance analysis of novel toxin-antidote CRISPR gene drive systems. bioRxiv, 2019.

28. Champer J, Lee YL, Yang E, Liu C, Clark AG, Messer PW. A toxin-antidote CRISPR gene drive system for regional population modification. bioRxiv, 2019.

29. Oberhofer G, Ivy T, Hay BA. Cleave and Rescue, a novel selfish genetic element and general strategy for gene drive. Proc Natl Acad Sci, 201816928, 2019.

